# ANASFV: a workflow for ASFV whole genome sequencing, assembly, and evaluation

**DOI:** 10.1101/2024.07.08.602471

**Authors:** Ke Li, Xu Han, Yanwen Shao, Yiwen Zhang, Xiaomin Zhao, Eager Wayne Johnson, Runsheng Li

## Abstract

African Swine Fever Virus (ASFV) poses an ongoing threat with widespread outbreaks affecting both domestic and wild pig populations globally. Effective management of ASFV outbreaks necessitates a deep understanding of its genetic diversity and evolutionary dynamics. Despite the advantages of nanopore sequencing for genome analysis, its application to ASFV genomes encounters specific challenges, such as high host DNA contamination that reduces viral genome coverage and an inherently higher error rate producing small insertions and deletions (indels). Another notable issue is the lack of standardized methods for assessing the quality of ASFV genomes. Furthermore, an increasing number of recombinant isolates of genotype I and II, have been observed, further complicating the task of phylogenetic analysis. To overcome these obstacles, we developed the ANASFV (analysis of an ASFV) pipeline. The pipeline is used to solve four tasks: In the first task, the pipeline introduces an amplicon sequencing approach that significantly improves genomic coverage, enabling reliable genome assembly. The pipeline also introduce reference-aided polishing techniques to correct small indels caused by nanopore sequencing errors. Moving on to the second task, a genome quality assessment system was established to evaluate the completeness and accuracy of the assembled genomes. For the third task, a method to rapidly analyze whether an isolate is a recombinant between genotype I and II, as well as to determine the pattern of recombination, based on gene similarity. Lastly, a comprehensive phylogenetic analysis based on coding sequences (CDS) was conducted, allowing us to generate a refined phylogenetic tree that includes all known ASFV genomes. The ANASFV pipeline will facilitate ASFV full genome sequencing using the nanopore platform. The pipeline will also support robust downstream bioinformatic analyses to assess genome integrity and conduct detailed phylogenetic studies based on whole-genome data.

## Introduction

The African Swine Fever Virus (ASFV) is the causative agent of African Swine Fever (ASF), a severe and often fatal viral disease that affects domestic and wild pigs(1). Since the initial identification of ASF in Kenya in 1921, the disease has emerged in countries throughout sub-Saharan Africa. The first outbreak of ASF outside of Africa occurred in the late 1950s, affecting both Europe and South America. In 2007, a new wave of ASF emerged in Georgia and rapidly expanded to other European countries, resulting in significant economic consequences for the global domestic pig industry(2). In August 2018, ASFV was identified in the northeastern region of China(3). The ASF outbreak in China had a significant impact due to the country’s high share in the global pork market, as China is the largest pork producer. By 2019, ASFV had spread rapidly throughout every region of China, causing widespread outbreaks and considerable economic losses in the domestic pig industry(4). The virus also extended its reach to other parts of East and Southeast Asia(5, 6). The spread of ASFV demonstrates the ongoing and far-reaching impact of ASFV, highlighting the urgent need for a global understanding of this virus.

ASFV is a virus with a genome size of approximately 180 Kb. Traditional short reads generated by Next-generation sequencing (NGS) platforms often struggle to accurately assemble the entire genome of ASFV due to its repetitive regions. In contrast, long reads produced by Oxford Nanopore Technologies (ONT) sequencing platforms are better suited for obtaining high-quality whole genome assemblies of ASFV. The increased read length provided by ONT technology enables the resolution of repetitive regions and facilitates the detection of structural variations within the viral genome, leading to more comprehensive and accurate genomic reconstructions. As of July 2024, 24 of the 406 ASFV isolates deposited in Genbank were assembled with the help of ONT sequencing. ONT-based sequencing methods offer rapid turnaround times compared to traditional sequencing techniques(7). The streamlined workflow, simplified library preparation, and real-time data analysis allow for faster generation of whole genome sequences. This speed is especially important during disease outbreaks or urgent research situations, where timely information is crucial for effective response and control measures.

Currently, there are limitations in the ONT sequencing of ASFV. First, the presence of substantial host DNA contamination poses a challenge in ONT sequencing, resulting in inadequate coverage of ASFV. Insufficient coverage leads to a fragmented and incomplete genome assembly. Second, ONT sequencing, especially with the R9.4.1 flow cell, exhibits a higher error rate, leading to the generation of numerous small indels(8). The indels generated are often found within consecutive identical bases (homopolymers) because of the challenge of accurate length identification of homopolymers by ONT. The presence of these indels can lead to fragmented genes, which can impact subsequent analyzes. In order to solve the error rate problem, researchers have used NGS-assistant polish(9), but this approach requires additional NGS reads.

The recombinant ASFV isolates of genotype I and II pose a significant threat due to their high lethality and transmissibility among pigs(10). The combine genetic elements from both genotype I and II, resulting in a mosaic genome that complicates disease control efforts and vaccine development. The live attenuated vaccines derived from genotype II ASFV have proven ineffective against these recombinant strains(10), underscoring the urgent need for new strategies to combat their spread. Since the first report in 2023, recombinant ASFV have been increasingly detected. Eleven isolates in NCBI are recombinants as of July 2024, originating from China, Russia, and Vietnam. Recombinant genotypes are a challenge for phylogenetic analysis. There is currently an ASFV biotyping tool based on p72(11). However, determining whether an isolate is recombinant requires the full genome of the isolate, and there is currently a lack of convenient tools for detecting recombination in ASFV.

Using whole CDS to build a phylogenetic tree is an ideal method to study the evolutionary relationships of ASFV isolates. ASFV has been classified into 24 different genotypes (I to XIV) based on sequence variations observed in p72(12). In Eurasia, the prevalent genotype II isolates show high similarity in their p72. So, differentiating genotype II isolates based on the p72 gene solely is insufficient(13, 14). The use of a full genome to build a phylogenetic tree also presents challenges due to the presence of numerous repetitive elements in the ASFV genome and difficulties in genome assembly for some isolates(10, 15, 16). Using only coding regions to build trees can avoid problems such as alignment problems in repeated regions and missing parts of the genome(17). Whole-proteome trees are more suitable for cross-species comparisons and are obviously not suitable when studying ASFV isolates. Therefore, the whole-CDS tree is a relatively ideal solution. Using whole CDS to build a phylogenetic tree can finely distinguish similar isolates and does not require highly intact genome assembly. As a result, the utilization of whole-CDS facilitates the incorporation of a greater number of isolates into the tree.

In early 2021, an outbreak of ASF occurred on a farm in Shandong, China, leading to significant livestock mortality. To investigate the outbreak, we developed a pipeline called ANASFV (analysis of an ASFV) that incorporates sequencing, assembly, polishing, completeness assessment, recombination test, and phylogenetic analysis of ASFV genomes. We incorporate PCR amplification with custom-designed primers to ensure adequate coverage. Moreover, we employed reference-guided polishing techniques to effectively rectify small indels introduced by nanopore reads, improving the accuracy of our analysis. These strategies overcome the limitations posed by the low amount of template DNA and the potential sequencing errors associated with ONT sequencing. And we developed a system for completeness evaluation and a tool for recombination test of the ASFV genome. We construct a phylogenetic tree of 297 ASFV isolates using 188 CDS. We tested this pipeline using the isolate ShanDong2021. As a result, we obtained a high-quality ShanDong2021 genome and a refined ASFV phylogenetic tree.

## Methods

### Sample Collection and Sequencing

The sample was collected from the lung tissue of sick pigs. Genomic DNA was extracted using the QIAamp MinElute Virus Spin Kit (Qiagen, Germany). Two sets of primers were designed with amplification product lengths of 20 Kb and 5 Kb (Table S1, Table S2). The samples were subjected to PCR amplification using the two primer sets, respectively. Subsequently, the amplified DNA products were prepared for sequencing following the nanopore sequencing library preparation protocol (LSK-110, ONT). The library was sequenced in R9.4.1 flow cells (FLO-MIN106D, ONT).

### Basecalling and Trimming

The raw nanopore sequencing data was basecalled by Guppy (v5.0.11) with the Super-accurate basecalling model. Subsequently, the reads were filtered and trimmed using NanoFilt(18) (v2.8.0), with a minimum average read quality score of 10 and a minimum read length of 1000 base pairs. Additionally, the first 50 bases were trimmed using the headcrop option.

### Genome Assembly

The genome exhibiting the highest read mapping coverage among all available ASFV genomes from NCBI is regarded as a reference genome (ANASFV: find_near_ref.py). This reference genome was then utilized for mapping assembly of the sequenced data. The alignment was performed using the minimap2(19) (v2.17) aligner with the -a option, and high quality trimmed reads as input. The alignment results were saved in SAM format. From the SAM file, a consensus sequence was generated using samtools(20) (v1.17). First, the SAM file was converted to the BAM format, excluding unmapped reads (-F 4). The resulting BAM file was then sorted using the samtools sort command. Finally, the consensus sequence was extracted from the sorted BAM file using the consensus command of samtools.

### Polishing the Consensus Sequence

The consensus sequence was further polished using two rounds of error correction. First, medaka (v1.11.3) (https://github.com/nanoporetech/medaka) was used. The high-quality trimmed reads and the initial consensus sequence were provided as input to the medaka_consensus program. Next, homopolish(21) (v0.4.1) was utilized to perform additional error correction. We performed nBLAST on the consensus sequence against the NCBI database, and the closest isolate we obtained was ASFV/pig/China/CAS19-01/2019 (GenBank: MN172368.1). We use this genome as a reference for homopolish. The homopolish algorithm used the R9.4.pkl model for correction.

### Phylogenetic analysis

The ASFV genomes used to build a phylogenetic tree were downloaded from NCBI, covering data up until July 2024 (Table S3). All 188 CDS from NC_044959.2 (ASFV Georgia 2007/1) were used as a reference as NC_044959.2 (ASFV Georgia 2007/1) is frequently used as a reference in other studies(22, 23). We employed Exonerate(24) (v2.4.0) to fetch the homology gene sequences from each strain’s whole genome sequence. The CDS of each gene will be used for tree construction. If no hit could be generated by Exonerate with default cutoff, this gene will not participate in the final tree building. The CDS results were then utilized as input for the tree construction of "de-novo" mode using uDance(25) (v1.6.3), which employs RAxML (v8) for maximum likelihood tree building. Subsequently, the generated tree served as a backbone for a second iteration of tree construction by adjusting the backbone option to "tree" in uDance, allowing adding the unplaceable nodes during "de-novo" tree building. For the total 406 isolates used in our analysis, 16 isolates had CDS sequenced identical with other isolates; One isolate (MT847622.1) could not find its optimal placement in the tree. These 17 isolates were excluded in the final tree. As a result, a total of 389 isolates were shown in the phylogenetic tree. Tree visualization was performed using iTOL(26) (v6.9).

## Results

To overcome the difficulties of nanopore sequencing of ASFV and conduct a more refined phylogenetic analysis of ASFV, we have developed a comprehensive pipeline called ‘ANASFV’ that includes sequencing, assembly, polishing, recombination test, quality evaluation, and phylogenetic analysis of the ASFV genome (Figure 1). We used the DNA sample from the ShanDong2021 isolate to test our pipeline. After performing multiple PCRs, we obtained the amplicon DNA from the ASFV DNA template. Subsequently, we conducted nanopore sequencing using the amplified DNA. The ONT reads were then subjected to mapping assembly to obtain a draft genome. To improve accuracy, we performed two rounds of polishing using medaka and homopolish. Finally, we obtained the polished genome, which was subjected to quality assessment, resulting in a BUSCO(27) (Benchmarking Universal Single-Copy Orthologs) -like evaluation. The ShanDong2021 is a genotype II ASFV and showing no evidence of recombination, according to recombination test. Additionally, we obtained sequence files for each CDS across genomes of ShanDong2021 and other isolates from NCBI (Table S3) by employing exonerate. Finally, we utilize uDance to construct a phylogenetic tree based on the obtained CDS data. For details on the code, see GitHub (https://github.com/lrslab/anasfv). Researchers conducting whole genome analysis of ASFV using nanopore sequencing can refer to this workflow. For researchers utilizing other sequencing platforms, the downstream analysis of this project, such as completeness evaluation, recombination test, and phylogenetic analysis, can be utilized.

**Figure 1.**
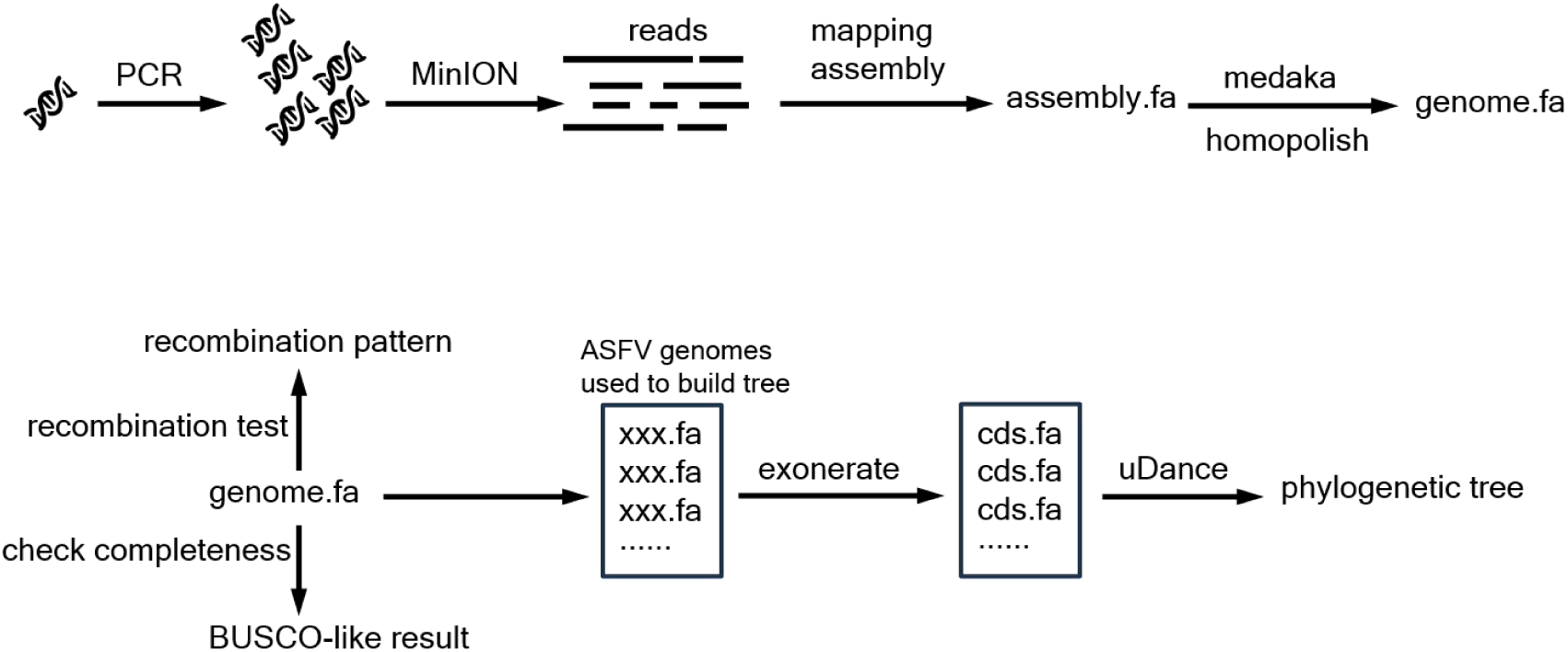
Workflow for ANASFV pipeline. Among all ASFV genomes from NCBI, the genome that ONT reads can map to the most will be used as the reference sequence for mapping assembly using homopolish.

### An amplicon system to amplify ASFV whole genome

Nanopore sequencing technology has good adaptability to processing long DNA fragments. We designed a set of primers with an amplification product of 20 Kb (Table S1), which is expected to make better use of the long-read characteristics of Nanopore sequencing technology and obtain more complete long-fragment sequencing data, but its amplification effect may not be good. Therefore, we also designed a set of primers with an amplification product of 5 Kb (Table S2). The 5 Kb set consists of 40 pairs of primers, with an overlap length between amplified products of approximately 100-500 bp. The 20 Kb set consists of 10 pairs of primers, with an overlap length between amplified products of approximately 500-900 bp. In this ShanDong2021 sample, the 5kb primer set designed for 5 Kb amplification product lengths demonstrated superior performance compared to the 20 Kb primer set in terms of amplification efficiency (Figure 2). There were 4 pairs of primers in the 20 Kb primer set that exhibited a suboptimal amplification efficiency. The 4 exceptions may have been influenced by factors such as primer design, template quality, or other experimental variables. The yield of reads for 20 Kb and 5 Kb was 748 Mb and 1.6 Gb, respectively (Table S4).

**Figure 2.**
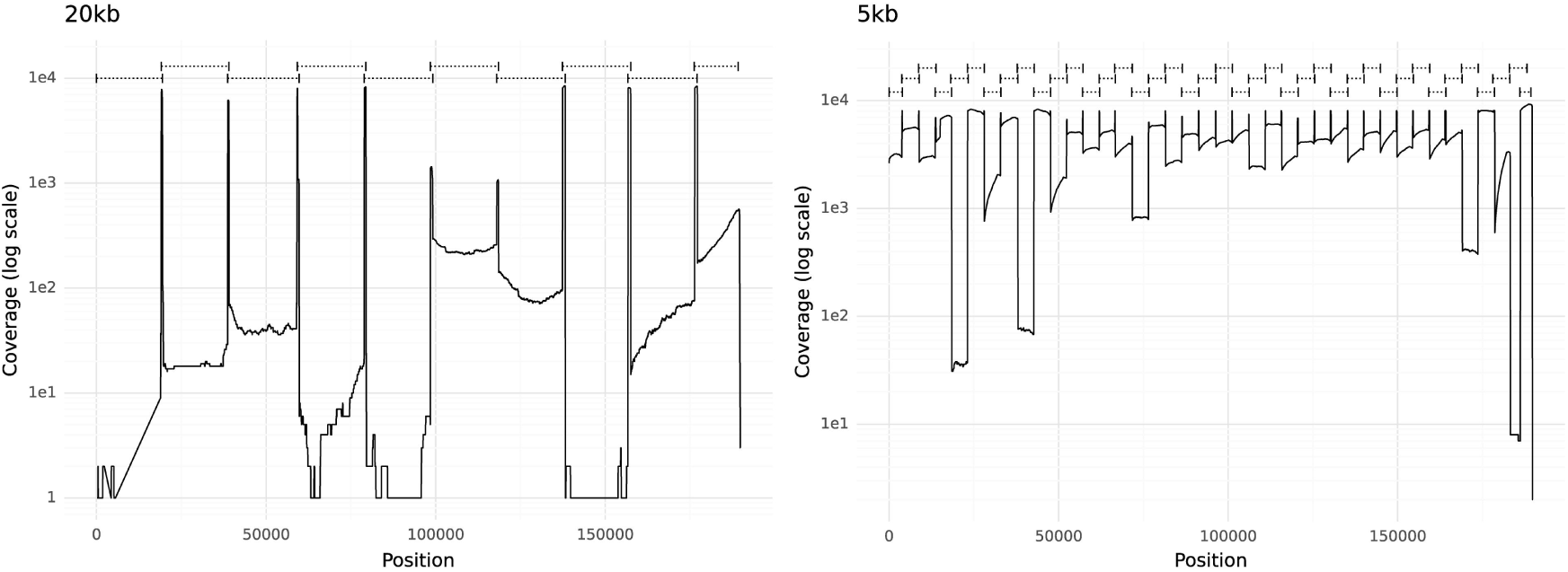
Coverage plot for ASFV amplicons. The logarithmic scale representation of the coverage depth across genomic positions of ShanDong2021, representing the results of amplification products sequencing from primer sets of 5 Kb (right) and 20 Kb (left), respectively. The positions of primer pairs used are indicated at the top of each figure.

### An evaluation system to evaluate the quality of the ASFV genome assembly

Currently, there is no standard for for assessing the completeness of ASFV genome. BUSCO is a widely used computational tool for assessing the completeness and quality of genome assemblies(28). Therefore, we developed a completeness evaluation system analogous to BUSCO. We generated consensus sequences for 148 CDS from genotype II isolates, with 33 MGF (multigene family) genes (MGF_100 : 2, MGF_110 : 3, MGF_300: 2, MGF_360 : 16, MGF_505 : 10), and consensus sequences for 149 CDS from genotype I isolates, with 34 MGF genes (MGF_100 : 2, MGF_110 : 4, MGF_300 : 2, MGF_360 : 16, MGF_505 : 10). The list of CDS used can be found in Table S5. CDS prediction of ShanDong2021 was performed using Prodigal(29) (v2.6.3), and the predicted CDS were compared to consensus sequences using BLASTN (e-value 1e-5). This comparison yielded a BUSCO-like genome completeness evaluation, with C:complete [D:duplicated], F:fragmented, M:missing, n:number of genes used. If a consensus CDS term can find a mapping from the predicted gene sequences with an identity larger than 90%, together with a unique mapping length longer than 90%, the consensus CDS term can be considered "complete." A consensus CDS term that cannot find a valid mapping (with an identity greater than 30% and a mapping length greater than 30%) in the predicted genes is considered "missing". The other consensus CDS with a partial hit is termed "fragmented". The "duplicated" term means that there is more than one ‘complete’ hit in the predicted gene sequences.

### Reference-aided polish helps to resolve the small indels generated by nanopore

The reads of amplicon sequencing provided sufficient coverage for accurate genome assembly, ensuring that no regions of the reference genomes were left unaddressed. We performed BLASTN on the draft assembly sequence against the NCBI database, and the closest isolate we obtained was ASFV/pig/China/CAS19-01/2019 (MN172368.1). We use this genome as a reference for downstream analysis. An evaluation of quality and accuracy was performed using QUAST (v5.2.0, default parameters) to compare our draft assembly with the reference genome. The evaluation revealed a mismatch rate of 7.98 per 100 Kb and an indel rate of 121.75 per 100 Kb. After performing reference-aided polishing using homopolish, the indel rate dropped to 1.6 per 100 Kb. Upon comparing the draft assembly with the consensus sequence of genotype II, we obtained the BUSCO-like notation of C: 52.7% [D: 0.0%], F: 45.95%, M: 1.35%, n: 148. After excluding the MGF gene, the notation was C: 54.78% [D: 0.0%], F: 43.48%, M: 1.74%, n: 115. Following the application of reference-aided polish using homopolish, we obtained a BUSCO-like notation of C: 99.32% [D: 0.0%], F: 0.68%, M: 0.0%, n: 148. And without MGF genes: C: 99.13% [D: 0.0%], F: 0.87%, M: 0.0%, n: 115.

### Completeness evaluation for all ASFV genomes in Genbank

We performed the BUSCO-like evaluation of all available isolates in NCBI as of July 2024 (Table S3). Among them, except ShanDong2021 (This study), 13 ASFV isolates only used ONT sequencing, as shown in Table 1. The completeness evaluation of these 13 isolates showed poor quality (complete < 94%). Among the poor-quality genomes, 12 isolates sequenced were clearly due to insufficient polish. The genomes of 12 isolates demonstrated a high gene count and significant gene fragmentation. Some coding regions in the ASFV genome (such as p72, p54 and p30, etc) are closely related to antigenicity. Sequencing errors in these regions can lead to indels, potentially resulting in inaccurate predictions of the antigenicity. Among the poor-quality genomes, three isolates have indels in the p72 (B646L) region, resulting in fragmented predictions of this gene. In BAN20221-4, at position 988 of the B646L, an deletion of a single "C" nucleotide has occurred, resulting in a truncation of the gene at this position. In ASFV/Kyiv/2016/131, at position 1138 of the B646L, an deletion of a single "T" nucleotide has occurred, resulting in a truncation of the gene at this position. In MSR2022S1, at position 1065 of the B646L, an deletion of a single "C" nucleotide has occurred, resulting in a truncation of the gene at this position. We applied reference-aided polish to the poor-quality genomes using the ANASFV pipeline. The gene fragmentation have been reduced after polishing (Figure 3A).

**Figure 3.**
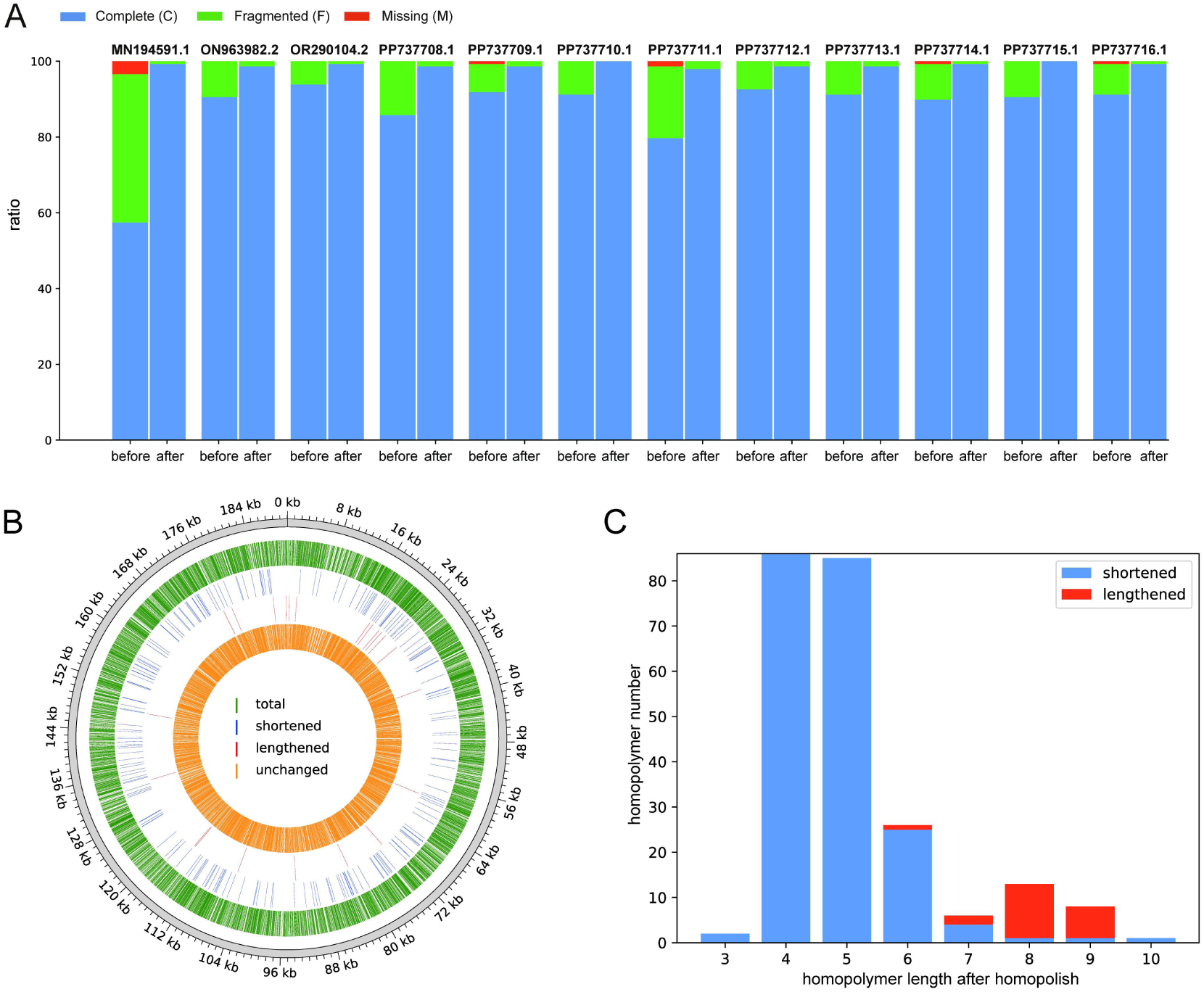
Changes in ASFV genome before and after homopolish. (A) Completeness evaluation of ASFV genomes that were sequenced by nanopore before and after homopolish. (B) The position for revised homopolymers on the ShanDong2021 genome. Only homopolymers with a length greater than three are shown. (C) The length distributions for revised homopolymers on ShanDong2021. Only homopolymers with lengths 3-10 are shown. Shortened, the homopolymer length becomes shorter after homopolish; Lengthened, the homopolymer length becomes longer after homopolish.

**Table 1.** Nanopore-based Genbank deposited ASFV whole genome sequences before and after polish.

|  |  |  |  |  |  |
| --- | --- | --- | --- | --- | --- |
| Accession ID | Isolate | Genome size<br>(before/after) | Prodigal gene num<br>(before/after) | Completeness(with MGF)<br>(before/after) | Completeness(without MGF)<br>(before/after) |
| MN194591.1 | ASFV/Kyiv/2016/131 | 191911/**191943** | 242/**169** | C:57.43%[D:0.0%],F:39.19%,M:3.38%,n:148<br>**C:99.32%[D:0.0%],F:0.68%,M:0.0%,n:148** | C:51.3%[D:0.0%],F:44.35%,M:4.35%,n:115<br>**C:100.0%[D:0.0%],F:0.0%,M:0.0%,n:115** |
| ON963982.2 | Philippines A4 2021 | 192265/**189530** | 186/**170** | C:90.54%[D:0.0%],F:9.46%,M:0.0%,n:148<br>**C:98.65%[D:0.0%],F:1.35%,M:0.0%,n:148** | C:91.3%[D:0.0%],F:8.7%,M:0.0%,n:115<br>**C:99.13%[D:0.0%],F:0.87%,M:0.0%,n:115** |
| OR290104.2 | CN/GD/2022 | 189416/**189443** | 174/**169** | C:93.92%[D:0.0%],F:6.08%,M:0.0%,n:148<br>**C:99.32%[D:0.0%],F:0.68%,M:0.0%,n:148** | C:93.91%[D:0.0%],F:6.09%,M:0.0%,n:115<br>**C:99.13%[D:0.0%],F:0.87%,M:0.0%,n:115** |
| OR449224.1 | Ratchaburi_2023_001-MA | 213885/**213883** | 217/**208** | C:91.89%[D:16.89%],F:0.68%,M:7.43%,n:148<br>**C:91.22%[D:18.92%],F:1.35%,M:7.43%,n:148** | C:97.39%[D:14.78%],F:0.0%,M:2.61%,n:115<br>**C:97.39%[D:14.78%],F:0.0%,M:2.61%,n:115** |
| OR660089.1 | ShanDong2021 | 190015/**189844** | 276/**169** | C:52.7% [D:0.0%], F:45.95%, M:1.35%, n:148<br>**C:99.32%[D:0.0%],F:0.68%,M:0.0%,n:148** | C:54.78% [D:0.0%], F:43.48%, M:1.74%, n:115<br>**C:99.13%[D:0.0%],F:0.87%,M:0.0%,n:115** |
| PP737708.1 | BAN20221-4 | 187609/**187654** | 192/**169** | C:85.81%[D:0.0%],F:14.19%,M:0.0%,n:148<br>**C:98.65%[D:0.0%],F:1.35%,M:0.0%,n:148** | C:86.96%[D:0.0%],F:13.04%,M:0.0%,n:115<br>**C:100.0%[D:0.0%],F:0.0%,M:0.0%,n:115** |
| PP737709.1 | PAN20211A | 189514/**189530** | 182/**170** | C:91.89%[D:0.0%],F:7.43%,M:0.68%,n:148<br>**C:98.65%[D:0.0%],F:1.35%,M:0.0%,n:148** | C:92.17%[D:0.0%],F:6.96%,M:0.87%,n:115<br>**C:99.13%[D:0.0%],F:0.87%,M:0.0%,n:115** |
| PP737710.1 | BTG2021KSU1-1 | 189540/**189550** | 182/**172** | C:91.22%[D:0.0%],F:8.78%,M:0.0%,n:148<br>**C:100.0%[D:0.0%],F:0.0%,M:0.0%,n:148** | C:91.3%[D:0.0%],F:8.7%,M:0.0%,n:115<br>**C:100.0%[D:0.0%],F:0.0%,M:0.0%,n:115** |
| PP737711.1 | MSR2022S1 | 189514/**189536** | 199/**169** | C:79.73%[D:0.0%],F:18.92%,M:1.35%,n:148<br>**C:97.97%[D:0.0%],F:2.03%,M:0.0%,n:148** | C:78.26%[D:0.0%],F:20.0%,M:1.74%,n:115<br>**C:98.26%[D:0.0%],F:1.74%,M:0.0%,n:115** |
| PP737712.1 | NEC20230726003 | 189537/**189547** | 183/**169** | C:92.57%[D:0.0%],F:7.43%,M:0.0%,n:148<br>**C:98.65%[D:0.0%],F:1.35%,M:0.0%,n:148** | C:93.04%[D:0.0%],F:6.96%,M:0.0%,n:115<br>**C:99.13%[D:0.0%],F:0.87%,M:0.0%,n:115** |
| PP737713.1 | NEC20230822001 | 189528/**189535** | 181/**170** | C:91.22%[D:0.0%],F:8.78%,M:0.0%,n:148<br>**C:98.65%[D:0.0%],F:1.35%,M:0.0%,n:148** | C:91.3%[D:0.0%],F:8.7%,M:0.0%,n:115<br>**C:99.13%[D:0.0%],F:0.87%,M:0.0%,n:115** |
| PP737714.1 | NEC20230929004A | 189539/**189543** | 180/**169** | C:89.86%[D:0.0%],F:9.46%,M:0.68%,n:148<br>**C:99.32%[D:0.0%],F:0.68%,M:0.0%,n:148** | C:89.57%[D:0.0%],F:9.57%,M:0.87%,n:115<br>**C:99.13%[D:0.0%],F:0.87%,M:0.0%,n:115** |
| PP737715.1 | NEC20230929004B | 189519/**189543** | 185/**169** | C:90.54%[D:0.0%],F:9.46%,M:0.0%,n:148<br>**C:100.0%[D:0.0%],F:0.0%,M:0.0%,n:148** | C:92.17%[D:0.0%],F:7.83%,M:0.0%,n:115<br>**C:100.0%[D:0.0%],F:0.0%,M:0.0%,n:115** |
| PP737716.1 | MDR202311F | 189501/**189514** | 183/**170** | C:91.22%[D:0.0%],F:8.11%,M:0.68%,n:148<br>**C:99.32%[D:0.0%],F:0.68%,M:0.0%,n:148** | C:90.43%[D:0.0%],F:8.7%,M:0.87%,n:115<br>**C:99.13%[D:0.0%],F:0.87%,M:0.0%,n:115** |

### Correction of homopolymers in the genome of ShanDong2021

A homopolymer refers to a polymer chain composed of the same type of nucleotide repeated consecutively, typically when a single nucleotide is repeated more than 2 times in a sequence. The ASFV genome contains an excessive number of homopolymers. We compared the number of homopolymers in the ASFV genome and human genome, together with the simulated random genome sequences with the same GC content of the ASFV (38.38%) or human genome (41%). The results show that the frequency of homopolymers with lengths greater than 3 in the ASFV genome is higher than in the human genome, and both are higher than in the simulated genomes (Figure S1). Although the total count of homopolymers in the ASFV genome is higher than in the human genome, homopolymers composed of "A" and "T" bases occur more frequently in the human genome compared to those of the ASFV. On the other hand, homopolymers composed of "G" and "C" bases occur with a higher frequency in the ASFV genome (Figure S2). There are 4866 homopolymers with lengths greater than 3 in the ShanDong2021 genome. After homopolish was applied, 229 homopolymers underwent changes. Among these 229 changes, 22 were lengthened, while the majority, specifically 207, were shortened (Figure 3C). In addition to the 229 typical homopolymer changes after homopolish, there was also one instance where "AA" was changed to "A" after homopolish. Among the indels generated by nanopore sequencing, 90.4 % (208 out of 230) were recognized as longer than their actual length and were thus shortened after homopolish. Most of these (198 out of 208) involved homopolymers ranging from 3 to 6 in length. The remaining 9.6% (22 out of 230) indels, which were recognized as shorter than their actual length, consisted of homopolymers ranging from 6 to 9 in length (Figure 3C).

### Description of Shandong2021

After polishing, we finally obtained an 189 Kb ASFV genome of ShanDong2021 (Accession ID: OR660089) with 72 genes in the forward strand and 97 genes in the reverse strand (Figure 4A). In the comparative analysis of the whole genome, ShanDong2021 exhibited an insertion of approximately 250 bp at the end of MGF_360-21R (Figure 4B), causing the gene length to increase from 1071 bp to 1092 bp. It can be seen from the dotplot (Figure 4B) that the repeat sequences (mainly MGF genes) are located at the beginning and the end of the genome, with more at the beginning than at the end.

**Figure 4.**
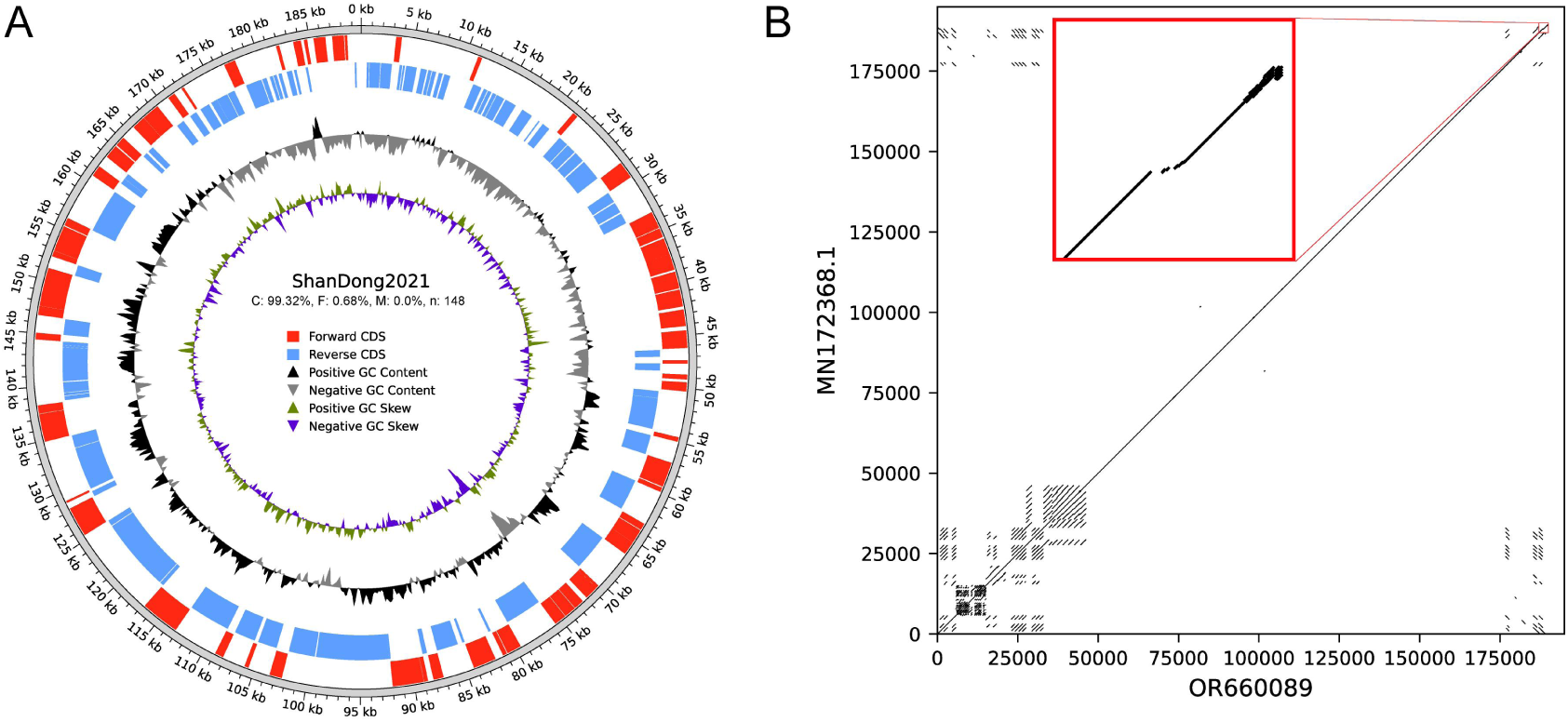
Genomic analysis of ShanDong2021. (A) Circular map of ShanDong2021 genome. (B) Dotplot of ShanDong2021 (OR660089) to ASFV/pig/China/CAS19-01/2019 (MN172368.1). The red box shows a zoomed-in view of the 187 Kb to 190 Kb region.

### A recombination test tool for checking recombinat of genotype I and II isolates

We generated consensus CDS from genotype I and II isolates. The list of CDS used can be found in Table S6. The CDS predicted from the ASFV genome using Prodigal were compared to the consensus sequences using BLASTN with an e-value threshold of 1e-5. For the alignment to be considered valid, the length of the alignment must be at least 80% of the consensus CDS length. If the identity to genotype I was higher, the CDS was classified as genotype I. If the identity to genotype II was higher, the CDS was classified as genotype II. If there was no BLASTN hit or the scores for both genotypes were equal, the CDS was marked as uncertain. We found significant differences in the B169L gene among the genotype I isolates (Figure 5A). The "GTCCAAAGCCGGCCG" motif appears once in B169L gene of genotype II ASFV, appears twice in tandem repeats in B169L gene of most genotype I ASFV, and appears three times in tandem repeats in the alternative B169L gene of genotype I ASFV. Five genotype I isolates (NHV, OURT 88/3, Pig/SD/DY-I/2021, Pig/HeN/ZZ-P1/2021, Benin 97/1) possess this alternative B169L gene, and all recombinant isolates also carry this alternative B169L gene. Therefore, in order to more accurately identify the gene, we also added the alternative B169L to the consensus CDS set of genotype I.

**Figure 5.**
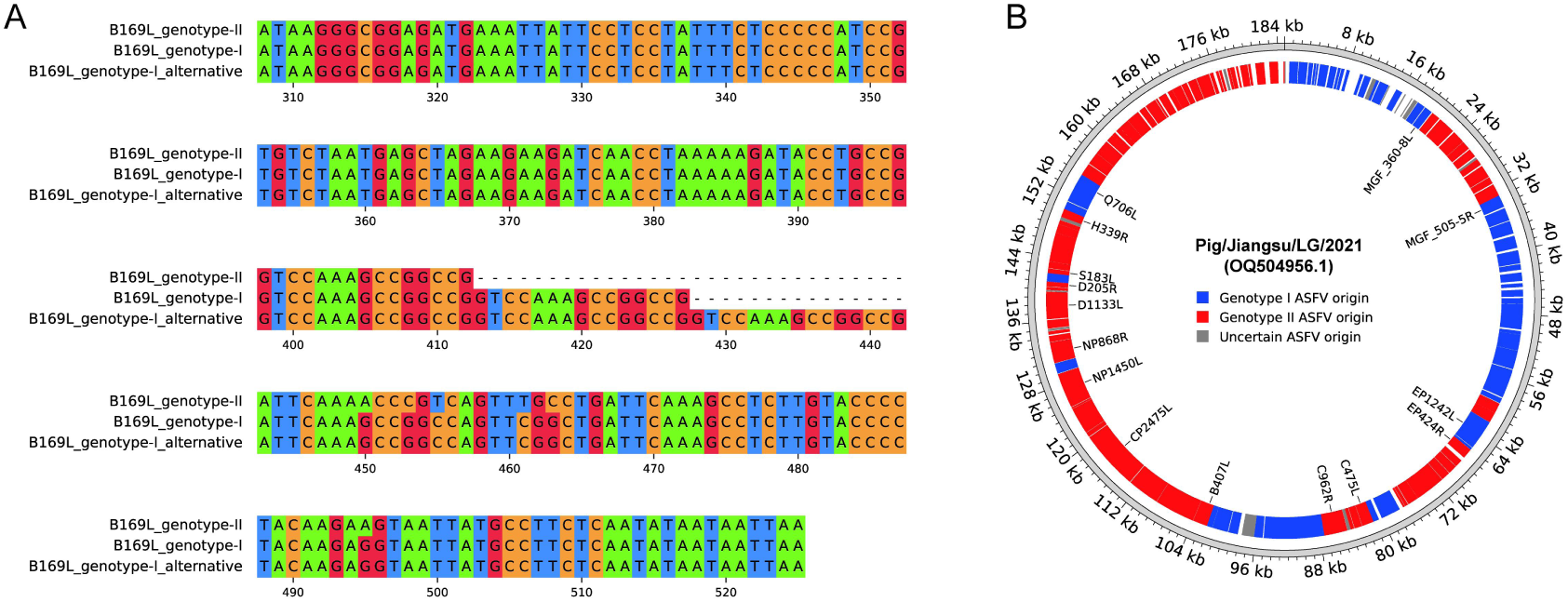
Recombination analysis. (A) An alternative B169L of genotype I ASFV compared with B169L gene consensus in genotype I and genotype II. (B) Recombination plot of a recombinant ASFV Pig/Jiangsu/LG/2021. The 15 genes with name labeled are genes that have a recombination within them.

### Recombination test for all ASFV genomes in Genbank

We performed the recombination test on all available isolates in NCBI as of July 2024. Eleven isolates were identified as recombinant of genotypes I and II (Table 2). The recombination patterns of a recombinant isolate Pig/Jiangsu/LG/2021 (Accession ID: OQ504956.1) were shown as an example (Figure 5B). The existing ASFV recombinants have 15 sub-gene recombination events. Since our method does not involve the sub-gene level, the recombination within genes is not reflected in the figure. The recombination patterns of all 11 recombinant isolates were consistent (Figure S3).

**Table 2.** Isolates that identified as recombinat of genotype I and II by ANASFV recombination test.

| Accession ID | Isolate | Country |
| --- | --- | --- |
| OQ504954.1 | Pig/Henan/123014/2022 | China |
| OQ504955.1 | Pig/Inner Mongolia/DQDM/2022 | China |
| OQ504956.1 | Pig/Jiangsu/LG/2021 | China |
| PP348677.1 | Primorsky 2023<br>DP-4560.Rec | Russia |
| PP478517.1 | NJS23-1 | China |
| PP478518.1 | NJS23-2 | China |
| PP712068.1 | JX23-01 | China |
| PP712069.1 | JX23-02 | China |
| PQ010732.1 | rASF12-avac02 | Vietnam |
| PQ010733.1 | rASF1/2-avac03 | Vietnam |
| PQ010734.1 | rASF1/2-avac07 | Vietnam |

### Phylogenetic analysis

We used all available ASFV genomes on NCBI as of July 2024 (Table S3) to make the phylogenetic tree, including 12 genotypes and recombinant type of I-II (Figure 6). Six distinct clades can be identified. Isolates VI, XX, IV, III, and XXII, which are considered to be different genotypes, are mixed in clade 3. Clade 3 also contains LIV_5/40(30), an isolate considered to be genotype I. However, this is not the first time this mixed clade has been noticed. Clade 3 is consistent with that of epsilon clade(31) and Biotype 3(22) in previous studies. There are 256 genotype II and 89 genotype I isolates in the tree. Eleven recombinants are located at the root of clade 1. Some genotype II isolates with poor genome quality are also located at the root of clade 1, such as ASF/IND/20/CAD/543 at the base (completeness: C:85.14%[D:0.0%], F:3.38%, M:11.49%, n:148). The ShanDong2021 isolate was clustered with LYG18 (OM105586) and ASFV-wbBS01 (MK645909), both of which were sampled from China. ShanDong2021 and ASFV-wbBS01 formed a small clade, indicating the closest evolutionary distance between the two. We identified three genes that showed significant divergence between the two genomes: I196L (Ka/Ks: 0.24238), M1249L (Ka/Ks: 0.335183), MGF_360-21R (Ka/Ks: 0.436954) (Table S8).

**Figure 6.**
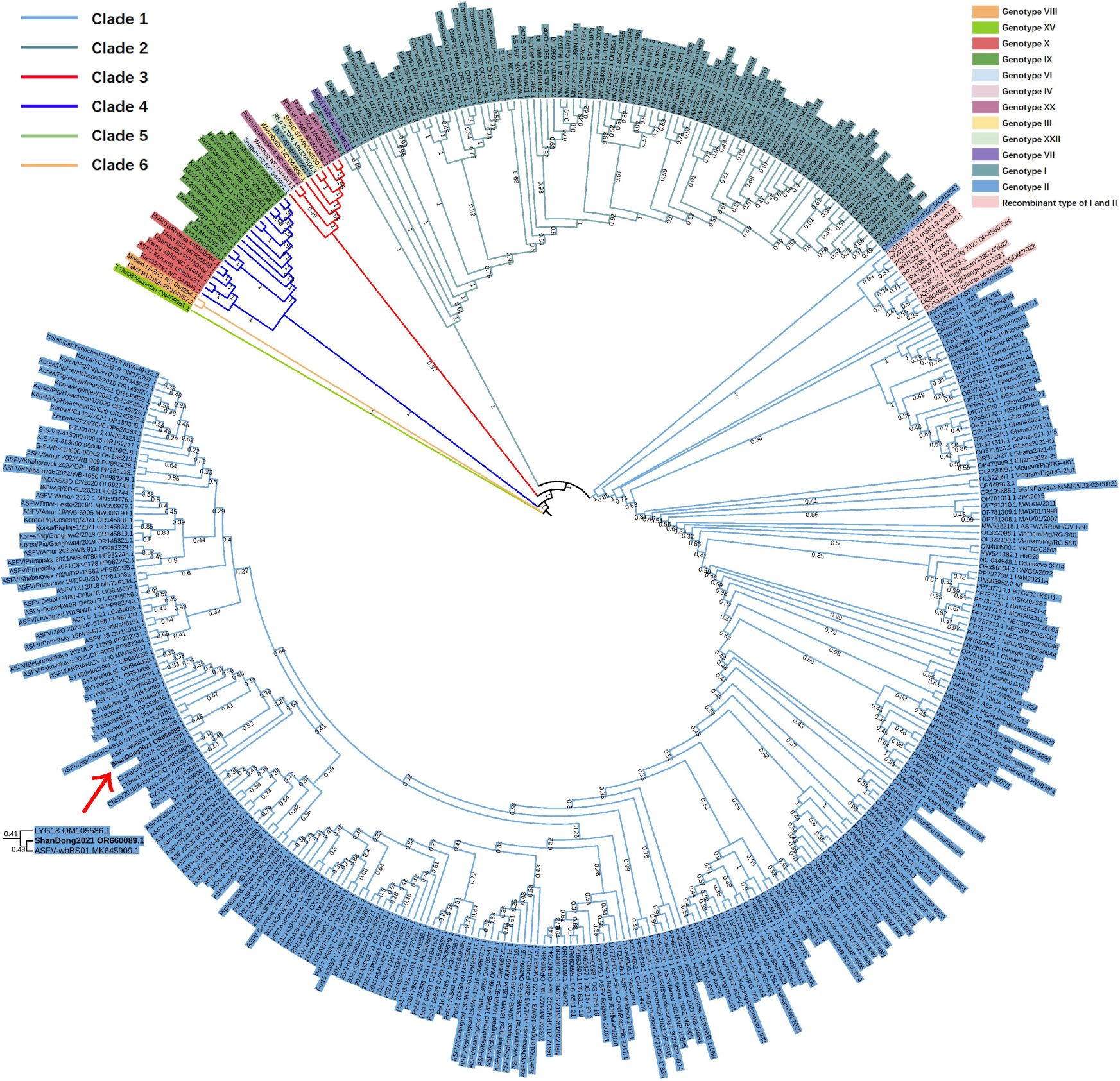
Phylogenetic analysis with all available ASFV whole genomes. The ShanDong2021 isolate is highlighted using bold letters and a red arrow. A magnified image of the branch of ShanDong2021 is shown in the lower right corner. Isolates are represented using their NCBI nucleotide accession numbers and their isolate names. The colors of the branches indicate the six clades identified by our study. The background color of the isolate name indicates different genotypes.

## Discussion

The read length of NGS is usually less than 500bp, which brings difficulties to genome assembly, especially repetitive regions. There are such repetitive regions at both ends of the ASFV genome, and genome assembly based on NGS reads cannot solve this problem well. The reads of ONT can reach ultra-long (longest >4 Mb), which can directly cover most repetitive regions, making the assembled genome more consistent with the real situation. The feasibility and effectiveness of amplicon sequencing for the ASFV genome, using primers targeting 10 Kb amplicons, have been supported by a previous study(32). In our study, the primer set of 5 Kb amplicon outperformed the primer set of 20 Kb amplicon. However, this preference is not absolute, as the amplification efficiency depends on the quality of the DNA. For specific needs, researchers can utilize primerdiffer(33) to design their own primer set accordingly.

The development of the ANASFV completeness evaluation system for the ASFV genome has provided a valuable tool to assess the quality of genome assemblies. Comparing genes in the genome to known consensus gene sets allows for the assessment of gene completeness and the identification of missing, duplicated, or fragmented genes. The occurrence of these missing, duplication, or fragmentation issues generally implies poor genome assembly. Since there are relatively few available genomes for other ASFV genotypes, we only established consensus gene sets for genotype I and genotype II. Therefore, ANASFV completeness evaluation is more effective for the currently prevalent genotype I and genotype II. The ANASFV completeness evaluation pipeline fills a gap in the tools available, allowing researchers to conveniently assess the quality of the ASFV genome assembly.

In the genome polishing process, we employed reference-aided polish, which has shown remarkable effectiveness in reducing errors introduced during ONT sequencing. Currently, most of the ASFV genomes on NCBI that are based only on ONT sequencing have the problem of insufficient polish. The results obtained from applying the completeness evaluation system to the nanopore-based ASFV genomes demonstrated the effectiveness of reference-aided polish in improving ONT sequencing errors in the absence of NGS reads as a reference for polishing. For researchers who only use ONT sequencing to obtain the ASFV genome, it is beneficial to employ reference-aided polishing to address indel errors that may persist due to insufficient polish by Medaka.

In our study, uDance was used to construct the whole-CDS tree(25). The uDance implements a divide-and-conquer strategy that simplifies the phylogenetic tree construction process. By utilizing parallel processing, uDance significantly reduces the time needed for tree construction. The refined phylogenetic tree provided valuable information on the genetic diversity and relatedness of different ASFV isolates, contributing to a better understanding of the epidemiology and evolution of ASFV. The ANASFV pipeline automates the process of downloading all available ASFV genomes from the NCBI database, performing CDS prediction and alignment. In case starting from scratch is not desired, we also provide our tree file (newick format), which can serve as a backbone tree to add new isolates by adjusting the backbone option to "tree" in uDance.

In general, our study demonstrates the successful application of ONT sequencing coupled with custom-designed PCR primers for complete genome sequencing of the ShanDong2021 ASFV isolate. The use of homopolish further enhanced accuracy. The genome quality assessment system and recombinant test can give an evaluation of the assembled genome. The construction of a phylogenetic tree provided valuable insights into the genetic characteristics and evolutionary relationships of ASFV isolates. In December 2023, when the study was just started, there were only 4 ASFV genomes sequenced only with ONT and only 3 recombinant ASFV genomes on NCBI; by July 2024, there were 14 ASFV genomes sequenced only with ONT and 11 recombinant ASFV genomes on NCBI. This increase shows that more and more people are starting to use ONT ot sequence ASFV and more and more recombinant ASFV are appearing, demonstrating that the ANASFV pipeline has great potential to contribute to the field of ASFV research.

## Data Availability

Nanopore sequencing reads included in this study have been submitted to the Sequence Read Archive (SRA) under accession numbers SRR28789321 and SRR28789322. The whole genome assembly for our isolate has been deposited in GenBank under accession number OR660089.

## Conflicts of Interest

The authors declare that they have no conflicts of interest.

## Acknowledgments

This work was supported by the Hetao Shenzhen-Hong Kong Science and Technology Innovation Cooperation Zone Shenzhen Park Project (HZQB-KCZYZ-2021017); the Early Career Scheme from the Hong Kong Research Grant Council (project number CityU 21100521); the Hong Kong Health and Medical Research Fund (project number 08194126), and the New Research Initiatives Support from City University of Hong Kong (project number 9610497) to R.L.

We thank Xudong Liu from City University of Hong Kong for the homopolymer occurrence data of human genome.

## Supplementary Materials

Supplementary_figures.docx :

Figure S1: Frequency of homopolymer occurrence in ASFV and human genomes. Figure S2: Frequency of homopolymer occurrence in ASFV and human genomes whit four bases displayed separately. Figure S3: Reconbination plot of 11 recombinat ASFV.

Supplementary_tables.xlsx : Table S1: 20kb primer set. Table S2: 5kb primer set. Table S3: Completeness evaluation of all available ASFV genomes. Table S4: Statistics of reads from 20kb and 5kb primer set. Table S5: Consensus gene list for completeness evaluation. Table S6: Consensus gene list used for recombination test. Table S7: Homopolished sites of ShanDong2021. Table S8: The statistics of Ka/Ks between ShanDong2021 and ASFV-wbBS01

## Notes

https://trace.ncbi.nlm.nih.gov/Traces/?view=run_browser&acc=SRR28789321&display=metadata

https://trace.ncbi.nlm.nih.gov/Traces/?view=run_browser&acc=SRR28789322&display=metadata

https://www.ncbi.nlm.nih.gov/nuccore/OR660089.1/

### Competing Interest Statement

The authors have declared no competing interest.

### Summary of Updates

Added the recombination test section and adjusted the content. The ASFV data used was updated from December 2023 to July 2024

## References

1. Blome S, Franzke K, Beer M. African swine fever–A review of current knowledge. Virus research. 2020;287:198099.

2. Rowlands RJ, Michaud V, Heath L, Hutchings G, Oura C, Vosloo W, et al. African swine fever virus isolate, Georgia, 2007. Emerging infectious diseases. 2008;14(12):1870.

3. Ge S, Li J, Fan X, Liu F, Li L, Wang Q, et al. Molecular characterization of African swine fever virus, China, 2018. Emerging infectious diseases. 2018;24(11):2131.

4. Ito S, Bosch J, Martínez-Avilés M, Sánchez-Vizcaíno JM. The evolution of African swine fever in China: a global threat? Frontiers in veterinary science. 2022;9:828498.

5. Jo YS, Gortázar C. African swine fever in wild boar, South Korea, 2019. Transboundary and Emerging Diseases. 2020;67(5):1776–80.

6. Nga BTT, Tran Anh Dao B, Nguyen Thi L, Osaki M, Kawashima K, Song D, et al. Clinical and pathological study of the first outbreak cases of African swine fever in Vietnam, 2019. Frontiers in Veterinary Science. 2020;7:392.

7. Petersen LM, Martin IW, Moschetti WE, Kershaw CM, Tsongalis GJ. Third-generation sequencing in the clinical laboratory: exploring the advantages and challenges of nanopore sequencing. Journal of clinical microbiology. 2019;58(1):10.1128/jcm.01315-19.

8. Ni Y, Liu X, Simeneh ZM, Yang M, Li R. Benchmarking of Nanopore R10. 4 and R9. 4.1 flow cells in single-cell whole-genome amplification and whole-genome shotgun sequencing. Computational and Structural Biotechnology Journal. 2023;21:2352–64.

9. Chen Z, Erickson DL, Meng J. Polishing the Oxford Nanopore long-read assemblies of bacterial pathogens with Illumina short reads to improve genomic analyses. Genomics. 2021;113(3):1366–77.

10. Zhao D, Sun E, Huang L, Ding L, Zhu Y, Zhang J, et al. Highly lethal genotype I and II recombinant African swine fever viruses detected in pigs. Nature Communications. 2023;14(1):3096.

11. Dinhobl M, Spinard E, Birtley H, Tesler N, Borca MV, Gladue DP. African swine fever virus P72 genotyping tool. Microbiology Resource Announcements. 2024;13(2):e00891–23.

12. Qu H, Ge S, Zhang Y, Wu X, Wang Z. A systematic review of genotypes and serogroups of African swine fever virus. Virus Genes. 2022;58(2):77–87.

13. Ankhanbaatar U, Auer A, Ulziibat G, Settypalli TB, Gombo-Ochir D, Basan G, et al. Comparison of the Whole-Genome Sequence of the African Swine Fever Virus from a Mongolian Wild Boar with Genotype II Viruses from Asia and Europe. Pathogens. 2023;12(9):1143.

14. Wang X, Wang X, Zhang X, He S, Chen Y, Liu X, et al. Genetic characterization and variation of African swine fever virus China/GD/2019 strain in domestic pigs. Pathogens. 2022;11(1):97.

15. Hakizimana JN, Yona C, Makange MR, Kasisi EA, Netherton CL, Nauwynck H, et al. Complete genome analysis of African swine fever virus genotypes II, IX and XV from domestic pigs in Tanzania. Scientific Reports. 2023;13(1):5318.

16. Mazloum A, van Schalkwyk A, Shotin A, Igolkin A, Shevchenko I, Gruzdev KN, et al. Comparative analysis of full genome sequences of African swine fever virus isolates taken from wild boars in Russia in 2019. Pathogens. 2021;10(5):521.

17. Kapli P, Yang Z, Telford MJ. Phylogenetic tree building in the genomic age. Nature Reviews Genetics. 2020;21(7):428–44.

18. De Coster W, D’Hert S, Schultz DT, Cruts M, Van Broeckhoven C. NanoPack: visualizing and processing long-read sequencing data. Bioinformatics. 2018;34(15):2666–9.

19. Li H. Minimap2: pairwise alignment for nucleotide sequences. Bioinformatics. 2018;34(18):3094–100.

20. Li H, Handsaker B, Wysoker A, Fennell T, Ruan J, Homer N, et al. The Sequence Alignment/Map format and SAMtools. Bioinformatics. 2009;25(16):2078–9.

21. Huang Y-T, Liu P-Y, Shih P-W. Homopolish: a method for the removal of systematic errors in nanopore sequencing by homologous polishing. Genome biology. 2021;22(1):1–17.

22. Dinhobl M, Spinard E, Tesler N, Birtley H, Signore A, Ambagala A, et al. Reclassification of ASFV into 7 Biotypes Using Unsupervised Machine Learning. Viruses. 2023;16(1):67.

23. Kwon O-K, Kim D-W, Heo J-H, Kim J-Y, Nah J-J, Choi J-D, et al. Genomic Epidemiology of African Swine Fever Virus Identified in Domestic Pig Farms in South Korea during 2019–2021. Transboundary and Emerging Diseases. 2024;2024(1):9077791.

24. Slater GSC, Birney E. Automated generation of heuristics for biological sequence comparison. BMC bioinformatics. 2005;6:1–11.

25. Balaban M, Jiang Y, Zhu Q, McDonald D, Knight R, Mirarab S. Generation of accurate, expandable phylogenomic trees with uDance. Nature biotechnology. 2023:1–10.

26. Letunic I, Bork P. Interactive Tree Of Life (iTOL) v5: an online tool for phylogenetic tree display and annotation. Nucleic acids research. 2021;49(W1):W293–W6.

27. Simão FA, Waterhouse RM, Ioannidis P, Kriventseva EV, Zdobnov EM. BUSCO: assessing genome assembly and annotation completeness with single-copy orthologs. Bioinformatics. 2015;31(19):3210–2.

28. Seppey M, Manni M, Zdobnov EM. BUSCO: assessing genome assembly and annotation completeness. Gene prediction: methods and protocols. 2019:227–45.

29. Hyatt D, Chen G-L, LoCascio PF, Land ML, Larimer FW, Hauser LJ. Prodigal: prokaryotic gene recognition and translation initiation site identification. BMC bioinformatics. 2010;11:1–11.

30. Ndlovu S, Williamson A-L, Malesa R, Van Heerden J, Boshoff CI, Bastos AD, et al. Genome sequences of three African swine fever viruses of genotypes I, III, and XXII from South Africa and Zambia, isolated from ornithodoros soft ticks. Microbiology Resource Announcements. 2020;9(10):10.1128/mra.01376-19.

31. Aslanyan L, Avagyan H, Karalyan Z. Whole-genome-based phylogeny of African swine fever virus. Veterinary world. 2020;13(10):2118.

32. Johnston CM, Olesen AS, Lohse L, le Maire Madsen A, Bøtner A, Belsham GJ, et al. A Deep Sequencing Strategy for Investigation of Virus Variants Within African Swine Fever Virus-infected Pigs. Pathogens. 2024;13(2):154.

33. Ren X, Shao Y, Zhang Y, Ni Y, Bi Y, Li R. Primerdiffer: a python command-line module for large-scale primer design in haplotype genotyping. Bioinformatics. 2023;39(4):btad188.

