## Supplementary Figures for "ANASFV: a workflow for ASFV whole genome sequencing, assembly, and evaluation"

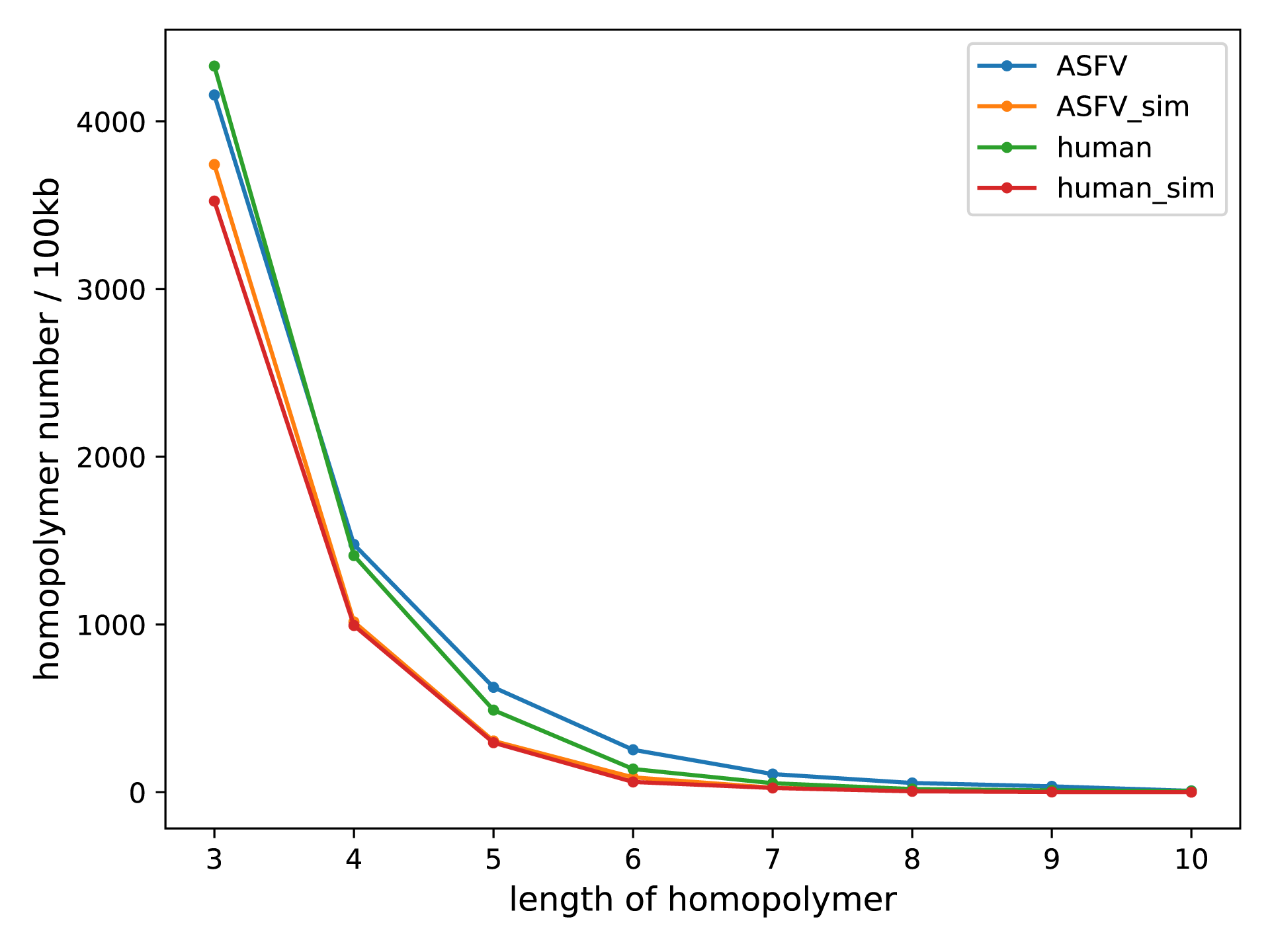
Figure S1. Frequency of homopolymer occurrence in ASFV and human genomes.“sim”, simulated genome using the same GC content.

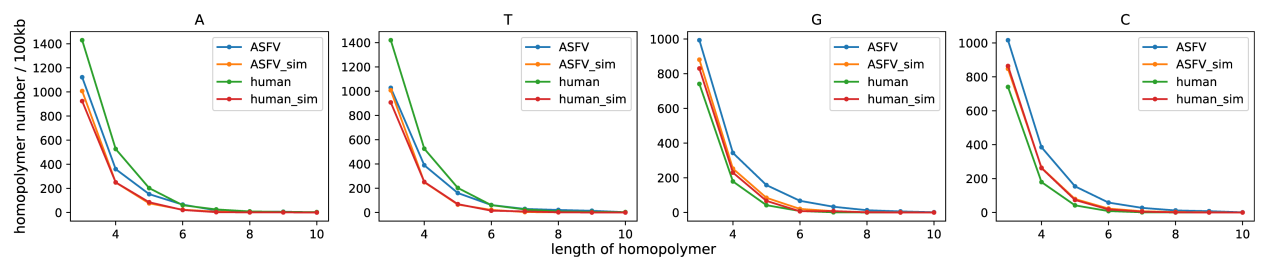

Figure S2. Frequency of homopolymer occurrence in ASFV and human genomes. The four bases are displayed separately.“sim”, simulated genome using the same GC content.

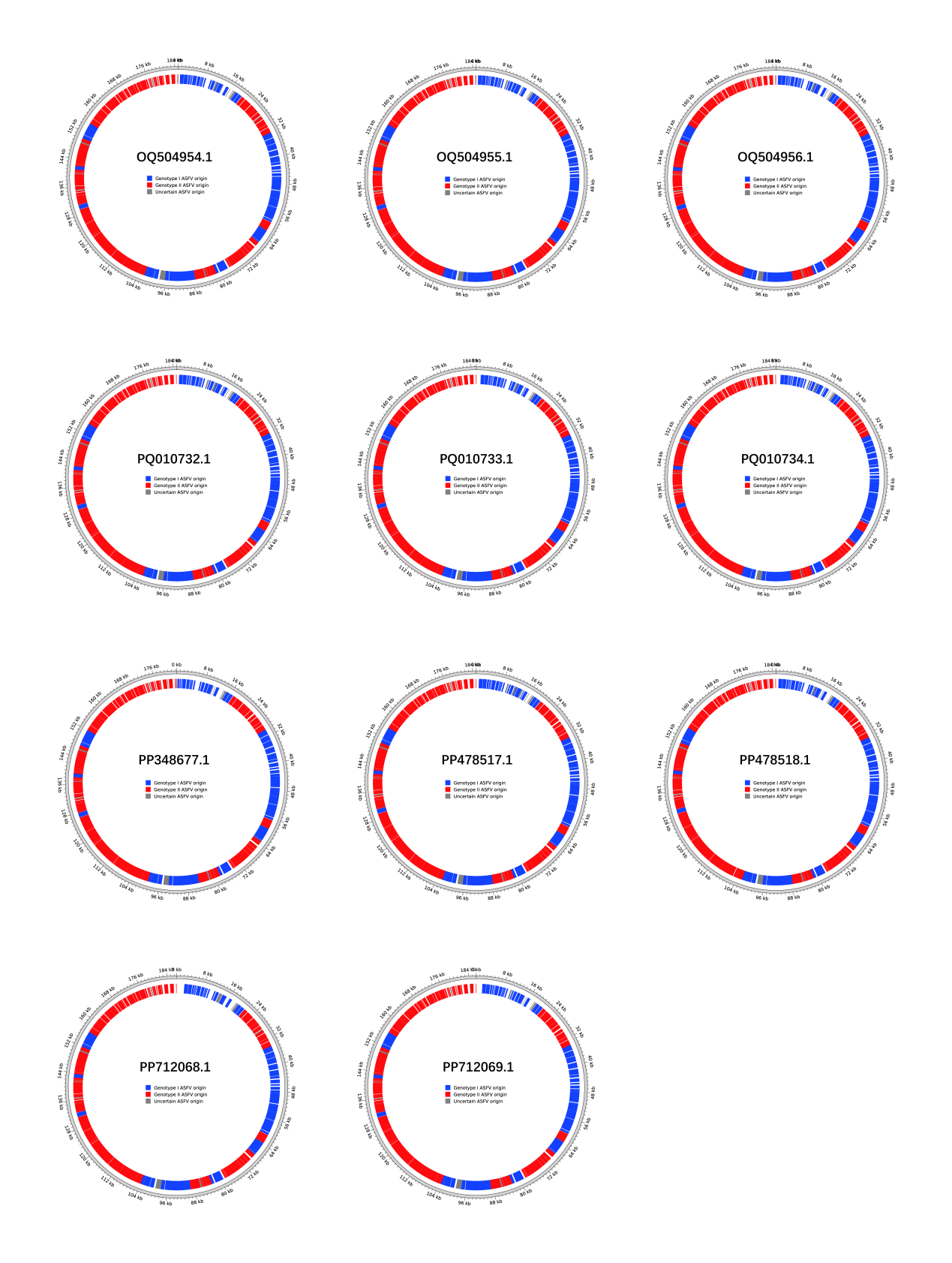
Figure S3. Reconbination plot of 11 recombinat ASFV isolates.
